# The ubiquitin ligase STUB1 suppresses tumorigenesis of renal cell carcinomas through regulating YTHDF1 stability

**DOI:** 10.1101/2023.08.23.554462

**Authors:** Siquan Ma, Yi Sun, Guoyao Gao, Jin Zeng, Ke Chen, Zhenyu Zhao

**Affiliations:** Department of Urology, Tongji Hospital, Tongji Medical College, Huazhong University of Science and Technology, Wuhan 430030, P.R.China; Hubei Institute of Urology, Wuhan 430030, P.R.China; Department of Urology, The First Affiliated Hospital of Nanchang University, Nanchang, Jiangxi 330000, P. R. China

**Keywords:** STUB1, YTHDF1, Ubiquitination, ccRCC, Cell migration, Cell invasion

## Abstract

STIP1 homology and U-box protein 1 (STUB1), a key RING family E3 ubiquitin ligase, plays both oncogenic and tumor-suppressive roles in a variety of human cancers. However, the role and mechanism of STUB1 in clear cell renal cell carcinoma (ccRCC) remains poorly defined. Here, we identified YTHDF1 as a novel STUB1 interactor by affinity purification mass spectrometry (AP-MS). STUB1 polyubiquitylates YTHDF1 and promotes YTHDF1 degradation. STUB1 depletion stabilizes YTHDF1 in renal cancer cells. STUB1-knockdown renal cancer cells exhibit increased migration and invasion in YTHDF1 dependent manner. Further study demonstrates that STUB1 knockdown promoted the tumorigenicity of ccRCC in a xenograft model. Clinically, STUB1 expression is down-regulated in ccRCC tissues, and the low expression level of STUB1 was associated with higher tumor stage and poor overall survival in patients with ccRCC. These findings reveal that STUB1 acts as an E3 ubiquitin ligase and promotes degradation of YTHDF1, and STUB1 inhibits the tumorigenicity of ccRCC through ubiquitinating YTHDF1.

**Novelty & Impact Statements:** STUB1 plays both oncogenic and tumor-suppressive roles in a variety of human cancers. Here, the authors demonstrated that STUB1 acts as a tumor suppressor in ccRCC, and the low expression level of STUB1 was associated with higher tumor stage and poor overall survival in patients with ccRCC. In addition, STUB1-knockdown renal cancer cells exhibit increased migration and invasion in YTHDF1 dependent manner. Mechanistically, STUB1 polyubiquitylates YTHDF1 and promotes YTHDF1 degradation.

## Introduction

Renal cell carcinoma (RCC) is a malignant tumor originating from the epithelium of the renal tubules; and the majority of RCC is clear cell renal cell carcinoma (ccRCC), which accounts for about 70-80% of RCC^[1]^. Overall, about 20-30% of RCC patients exhibit metastases at the time of initial diagnosis. In addition, about 40% of patients with localized RCC will develop distant metastases after primary surgery^[2]^. Currently, the treatments for metastatic ccRCC (mRCC) include anti-angiogenic drugs targeting VEGF and its receptors, mTOR inhibitors, and immune checkpoint inhibitors^[3]^. However, most of mRCC patients experience treatment failure after receiving the above treatments, and these patients are often lethal with a 5-year survival rate of only about 15%^[4]^. Thus, understanding the molecular mechanisms of ccRCC progression and metastasis is critical for identifying new drug targets for ccRCC patients.

STIP1 homology and U-box protein 1 (STUB1), also named as Carboxy-terminus of Hsc70/Hsp70-interacting protein (CHIP), is a RING family E3 ubiquitin ligase that acts along with Hsp90/Hsp70 chaperones and the ubiquitin-proteasome system (UPS) to regulate protein homeostasis. The best-known function of STUB1 is promotes the ubiquitylation and degradation of damaged proteins recognized by molecular chaperones. As an E3 ubiquitin ligase, STUB1 has s a wide range of substrate proteins and many of its substrates involve in regulating several physiological and pathological processes, such as cancer^[5]^. Consequently, STUB1 can regulate the occurrence and development of certain cancers. Indeed, increasing studies have demonstrated that STUB1 functions as a tumor-suppressive gene in multiple cancer types, including breast cancer^[6, 7]^, prostate cancer^[8-12]^, colorectal cancer^[13, 14]^, liver cancer^[15]^, lung cancer^[16, 17]^, pancreatic cancer^[18]^, ovarian cancer^[19]^, renal cancer^[20]^ and gastric cancer^[21]^. Importantly, Hsp90 inhibitors has anti-tumorigenic effect by promoting the STUB1 mediated ubiquitylation and degradation of specific oncoproteins^[5]^. Thus, overexpression of STUB1 expected to inhibit cancerous growth.

Interestingly, several studies indicate that STUB1 promotes cell proliferation and survival via degradation of its substrates such as PTEN^[12, 22]^, HIPK2^[23]^, PFN1^[24]^, p14ARF^[25]^, FOXO1^[26]^, and p21^[27]^. In addition, STUB1 was shown to be up-regulated in gallbladder carcinoma tissues when compared with their normal tissues, which was associated with a poor prognosis in gallbladder cancer^[28]^. Thus, STUB1 can play context-dependent functions in cancers. However, the role and the mechanism of STUB1 in ccRCC remains poorly defined.

In this study, we identified YTHDF1 as a novel STUB1 interactor by affinity purification mass spectrometry (AP-MS), and demonstrated that STUB1 as an E3 ubiquitin ligase for YTHDF1. We also showed that STUB1-knockdown renal cancer cells exhibit increased migration and invasion in YTHDF1 dependent manner. Moreover, we found that STUB1 knockdown promoted the tumorigenicity of ccRCC in a xenograft model. Importantly, STUB1 expression is down-regulated in ccRCC tissues, and the low expression level of STUB1 was associated with higher tumor stage and poor overall survival in patients with ccRCC. Overall, these results indicate that STUB1 suppresses tumorigenesis of renal cell carcinomas through regulating YTHDF1 stability.

## Results

### STUB1 expression in ccRCC tissues

Previous study has reported that STUB1 is down-regulated in renal cancer tissues^[20]^. To further confirm this result, we performed analysis of STUB1 protein expression in two publicly available datasets CPTAC-ccRCC (n = 110) and Chinese ccRCC (n=232) cohorts^[29, 30]^. As expected, this analysis revealed that the STUB1 protein level was significantly decreased in ccRCC tissues than in paraLcarcinoma non-tumor tissues in both the CPTAC and Chinese cohorts (Fig. 1A, B). Furthermore, we also performed immunohistochemical analysis of 12 paired ccRCC samples using a STUB1 antibody. The results indicated that STUB1 exhibits low expression levels in the tumor tissues compared to control samples (Fig. 1C, D, and Supplementary Figure S1). The above results further confirmed that STUB1 protein is downregulated in ccRCC.

**Figure 1.**
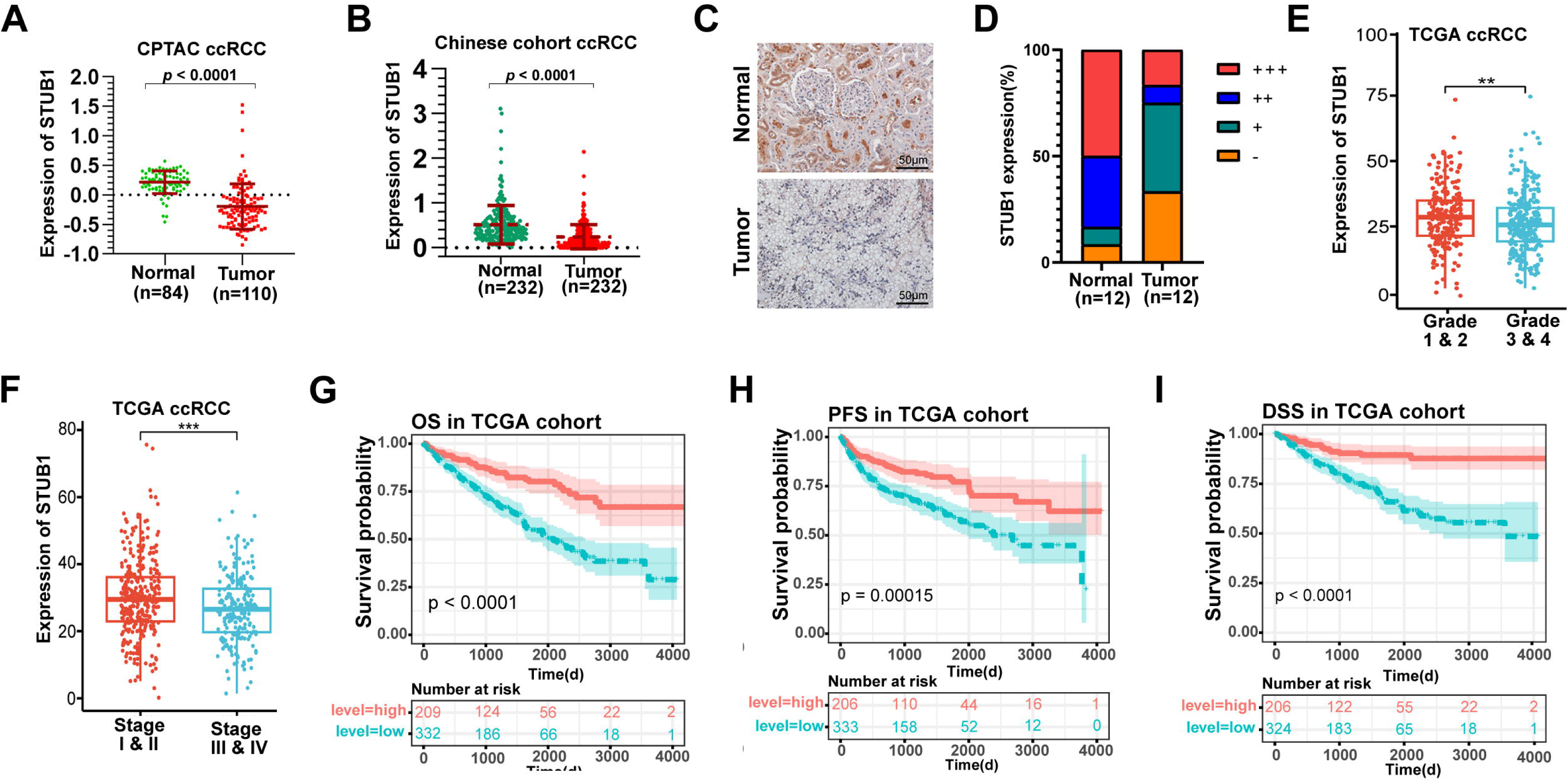
STUB1 expression in ccRCC. (**A**) The protein level of STUB1 between primary tumor tissues and non-tumor adjcant tissues of ccRCC in CPTAC cohort. (**B**) The protein level of STUB1 between primary tumor tissue and non-tumor adjcant tissues of ccRCC in 232 paired Chinese ccRCC cohort. (**C**) Immunohistochemical staining results of tumor and adjacent normal tissues of ccRCC patients. The magnification of the picture is 200 times, and the scale bar is 50 μm. (**D**) According to the score of immunohistochemical score system, the sections of ccRCC patients were divided into four grades: strong positive (+++), positive (++), weak positive (+) and negative (-). (**E**) Based on the TCGA data, the expression level of STUB1 were analyzed by the pathological grades of ccRCC. (**F**) The expression level of STUB1 in stage I-II and stage III-IV was compared according to the pathological stages. (**G-I**). Correlation between STUB1 gene expression and survival prognosis of ccRCC patients. Based on the TCGA data, low expression of STUB1 is associated with poor prognosis in patients with ccRCC. OS: Overall survival, PFS: Progression-free survival, DSS: Disease-special survival. ** *p*<0.01, *** *p*<0.001.

We further performed a comprehensive analysis of ccRCC datasets from the TCGA (The Cancer Genome Atlas) database and found that STUB1 expression was lower in tissues with higher histologic stage and grade tissues (Fig. 1E, F). In addition, the survival analysis revealed that low expression of STUB1 indicated a shorter overall survival (OS), progression-free survival (PFS) and disease-specific survival (DSS) (Fig. 1G-I). Together, these data suggested that the down-regulation of STUB1 may play an important role in renal cancer development.

### STUB1 is a tumor suppressor in ccRCC

To explore the function of STUB1 in ccRCC tumorigenesis, we established stably transfected cell lines expressing small hairpin RNA (shRNA) using lentivirus. As shown in Figure 2A and 2B, expression of two different STUB1 shRNAs can effectively down-regulated the mRNA and protein expression of STUB1 in 786-O and OS-RC-2 cells. Firstly, MTS and colony formation assays were performed to evaluate cell proliferation in these cells. Unexpectedly, the results showed that STUB1 depletion has little effect on the proliferation of 786-O and OS-RC2 cells (Fig. 2C-E). However, we observed that knockdown of STUB1 markedly increased both the migration and invasion capability of 786-O and OS-RC2 cells (Fig. 2F). The above evidence suggests that STUB1 is likely to function as a tumor suppressor and inhibit the tumorigenesis of ccRCC.

**Figure 2.**
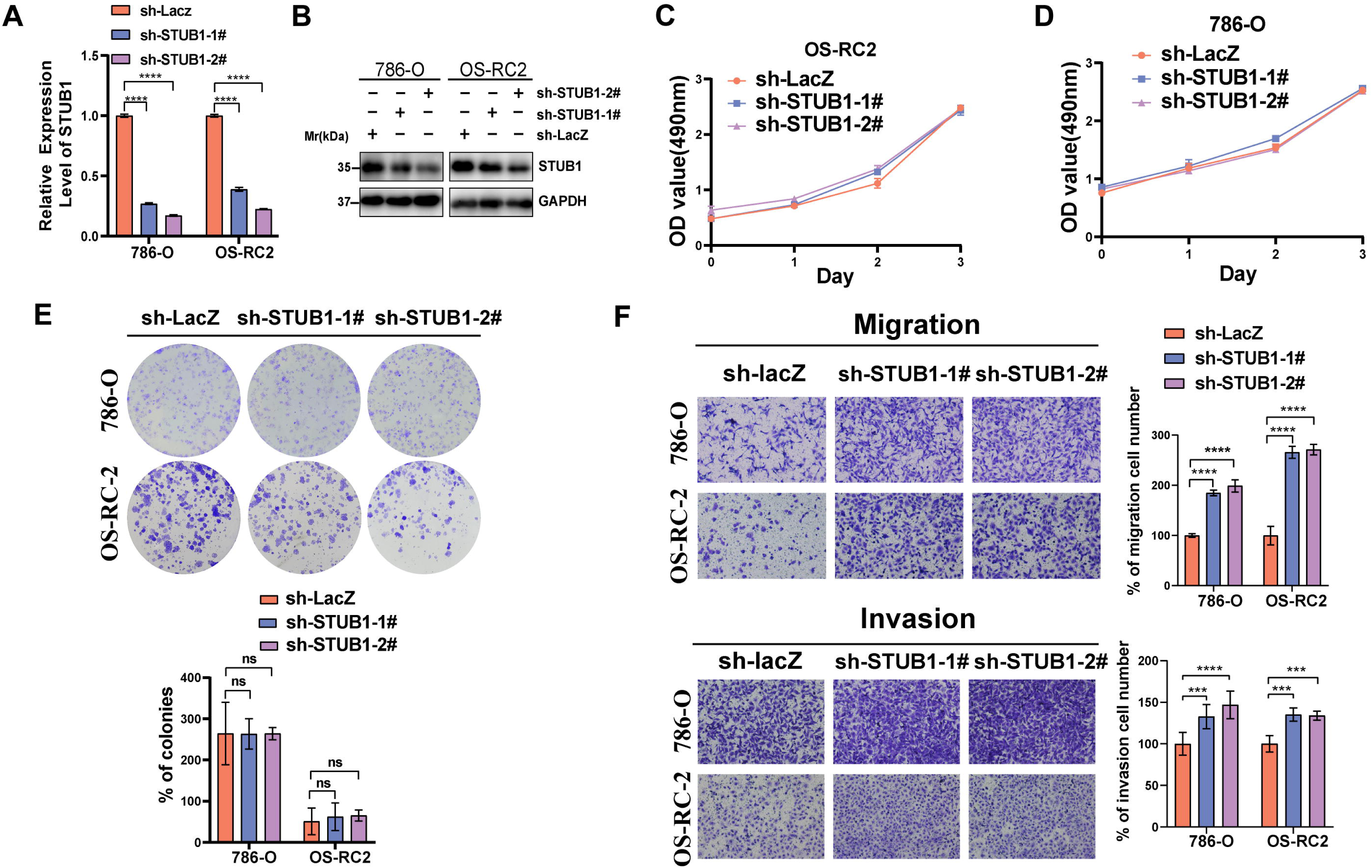
STUB1 knockdown promotes the cell migration and invasion of renal cancer cells. (**A**) qPCR was used to verify the knockdown efficiency of STUB1 in 786-O and OS-RC-2 cell lines. (**B**) . Immunoblotting was performed to evaluate the expression of STUB1 in 786-O and OS-RC-2 cells stably expressing sh-STUB1 or sh-LacZ. (**C, D**) MTS method was utilized to evaluate whether the knockdown of STUB1 affects the cell proliferation ability of 786-O and OS-RC-2 compared with the control group. (**E**) The 786-O and OS-RC-2 cells stably expressing sh-STUB1 or sh-LacZ and analyzed by colony formation assays. (**F**) Migration and invasion assays for 786-O and OS-RC-2 cells stably expressing sh-STUB1 or sh-LacZ. Migrated and invaded cells from each treatment group were counted in four random images. \**p*<0.05, \*\**p*<0.01, \*\*\**p*<0.001, \*\*\*\**p*<0.0001.

To explore whether STUB1 knockdown affects tumor growth *in vivo*, we established OS-RC-2 cells stably expressing doxycycline (dox)-inducible shRNA targeting STUB1 or control shRNA targeting LacZ (Fig. 3A), and then implanted these engineered OS-RC-2 cell lines into nude mice. After 7 days of implantation, dox was used to induce the expression of shRNA. Inconsistent with *in vitro* results, OS-RC2 xenografts samples showed a significant increase of tumor growth upon STUB1 shRNA induction (Fig. 3B-D). In addition, the number of blood vessels shown by CD31 immunohistochemical staining was significantly increased in STUB1-shRNA OS-RC-2 tumors as compared with the control tumors (Fig. 3E). Meanwhile, the expression of Ki-67 was also increased upon STUB1 shRNA induction (Fig. 3E). These data indicate that STUB1 acts as a tumor suppressor to inhibit renal cancer cell growth in mice.

**Figure 3.**
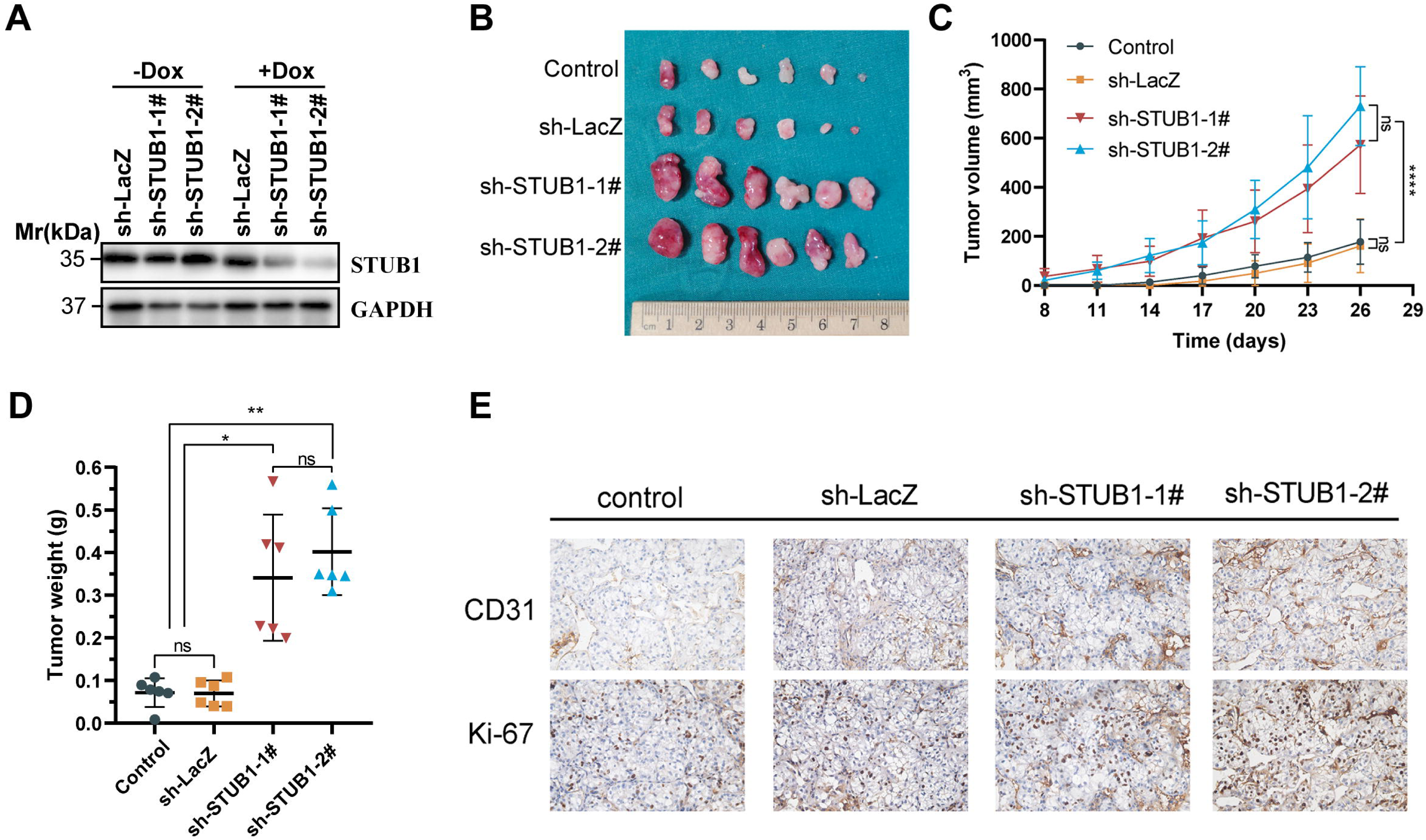
STUB1 konckdown promotes the RCC tumor growth *in vivo*. (**A**) OS-RC-2 cells expressing dox-inducible shRNAs were treated by dox. 72 h after dox treatment, the whole cell lysate was extracted for immunoblotting with anti-STUB1. (**B-D**) OS-RC-2 cells stably expressing dox-inducible sh-STUB1-1# and sh-LacZ (control) were subcutaneously injected into nude mice as indicated and dox was administered as described in the Method section. Tumor growth was measured every 3 days. Images (B), tumor growth curves (C) and weight (D) of xenografts are shown. (**E**) Representative immunohistochemical staining of CD31 and Ki-67 in xenograft tumors. \**p*<0.05, \*\**p*<0.01, ****p<0.0001.

### Identification of YTHDF1 as a noval STUB1 associated protein

To investigate the mechanism behind STUB1-mediated function in ccRCC, we conducted an affinity purification coupled MS (AP-MS) to identify the STUB1-interacting proteins. First, we established OS-RC-2 cells stably expressing StrepII-tagged STUB1, and then STUB1 was affinity-purification using Strep-Tactin beads and resolved by SDS-PAGE (Fig. 4A). After the bands in the SDS gel were analyzed by LC-MS/MS, we selected protein satisfying the following conditions as candidate interactions (Tables S1). (i) Two or more unique peptides corresponding to the protein were detected, and (ii) the number of the unique peptides in the Strep-II-STUB1 sample was > 2-fold larger than that in the empty vector control. Next, we queried the gene ontology cellular component (GO-CC) term association of these interactors and found an enrichment for proteins involved in the ‘nuclear speck’ (GO:0016607), ‘condensin complex’ (GO:0000796), ‘RNA polymerase II, holoenzyme’ (GO:0016591), ‘inclusion body’ (GO:0016234), ‘stress granule’ (GO:0010494), ‘chaperone complex’ (GO:0101031), ‘proteasome complex’ (GO:0000502), and ‘IkappaB kinase complex’ (GO:0008385) (Fig. 4B). We also performed a functional network analysis to identify possible complex formation between STUB1 interactions. The result showed that the 246 interactors were grouped into 9 subnetworks, and the largest subnetwork containing 181 seeds in which the central nodes are RPS27A, HSP90AA1, PCNA, DHX15, and POLR2E (Fig. 4C). In addition to heat shock proteins (HSPA2, HSPA1A, HSP90AB2P, HSP90AB1, HSP90AA1, HSPH1, HSPBP1, HSPA4L, and HSPA4) and proteasome complexes (PSMC5, PSMD6, PSMD11, PSMC6, PSMA5, ECPAS), we also identified some previously unreported proteins, including YTHDF1.

**Figure 4.**
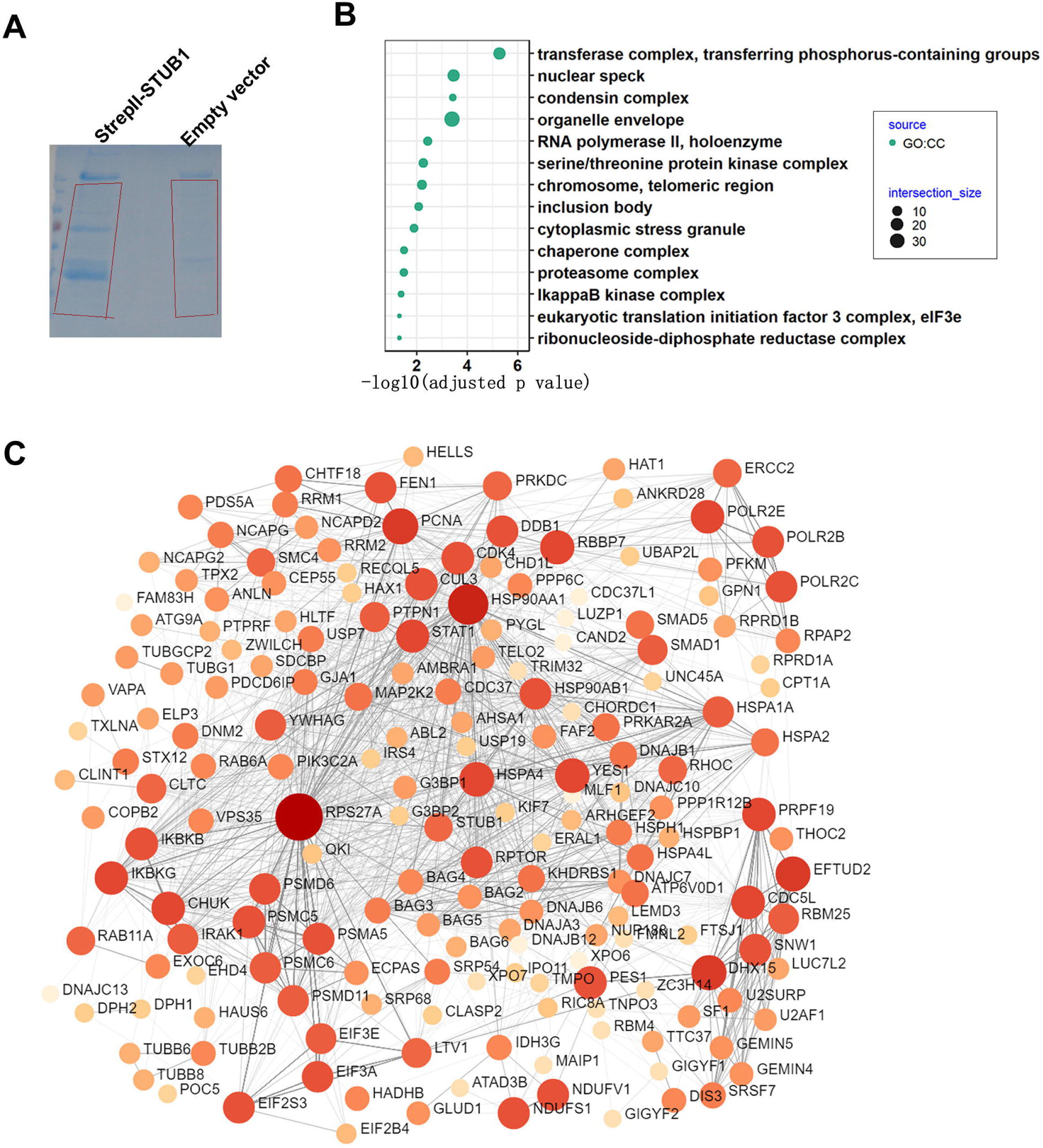
Identification and analysis of STUB1 interacting proteins. (**A**) Coomassie brilliant blue staining of StrepII-STUB1 precipitates resolved by SDS-PAGE. (**B**) Gene Ontology (GO) analysis of the 246 proteins identified as STUB1 partners in the HEK293T cells. (**C**) Network analysis of the 246 proteins identified as STUB1 partners.

We next performed co-precipitation experiment to prove the interaction between STUB1 and YTHDF1. HA-YTHDF1 was found in StrepII-STUB1 precipitates from HEK293T cells co-transfected with StrepII-STUB1 and HA-YTHDF1 plasmids (Fig. 5A). Reciprocal Co-IP assays were also performed in HEK293T cells expressing Myc-STUB1 and Flag-YTHDF1, Flag-YTHDF2, or Flag-YTHDF3, and the results showed that both YTHDF1 and YTHDF2 can pull down Myc-STUB1 (Fig. 5B). In order to eliminate any artifact due to excessive expression of proteins by transfection, we established HEK293T cells containing 2xFlag and Cherry at the C terminus of endogenous YTHDF1 (YTHDF1^Flag-Cherry^), and then we conducted Co-IP assay using anti-Flag antibody and found endogenous STUB1 was pulled down in the 2xFlag and Cherry knock-in cells (Fig. 5C). Additionally, to verify whether YTHDF1 and STUB1 directly interact, we added bacterially expressed GST-YTHDF1 into the lysates of HEK293T cells; and GST pull down assay showed GST-YTHDF1, but not GST protein, precipitated STUB1 (Fig. 5D). Last, Cherry-STUB1 co-localization with EGFP-YTHDF1 in the cytoplasm as revealed by fluorescent microscopy analysis (Fig. 5E). Together, the above results indicate that STUB1 directly interacts with YTHDF1 in human cells.

**Figure 5.**
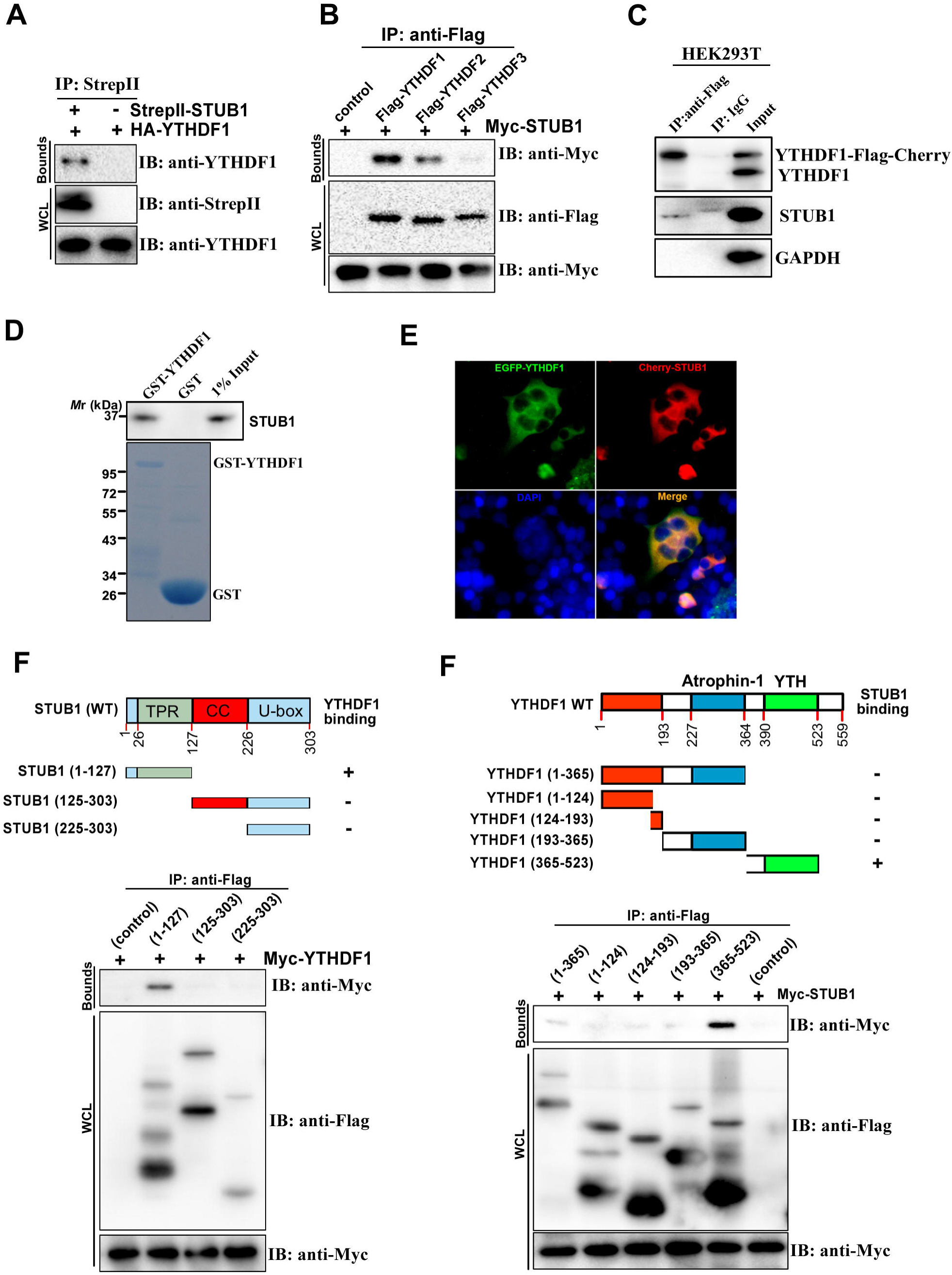
STUB1 interacts with YTHDF1. (**A**) Immunoblots of whole cell lysates (WCL) and immunoprecipitations (IP) from HEK293T cells co-transfected with StrepII-STUB1 and HA-YTHDF1 as indicated. (**B**) Immunoblots of WCL and IP from HEK293T cells transfected with Myc-STUB1 alone or along with Flag-YTHDF1, Flag-YTHDF2, Flag-YTHDF3 as indicated. (**C**) Co-IP assays of endogenous STUB1 and YTHDF1 (YTHDF1^Flag-Cherry^) in the 2xFlag and Cherry knock-in cells. (**D**) Pulled-down STUB1 by GST-YTHDF1 were detected by immunoblots with STUB1 antibody. Purified GST or GST-YTHDF1 were visualized by Coomassie brilliant blue staining. (**E**) Co-localization of STUB1 with YTHDF1 in the cytosol imaged by fluorescence microscopy. OS-RC-2 cells were co-transfected with Cherry-STUB1 and EGFP-YTHDF1. The cell nuclei were stained with DAPI. (**F**) Mapping of the STUB1 domain binding to YTHDF1. Schematic depiction of domain-deleted STUB1 (upper). Co-IP assays of Myc-YTHDF1 and truncated Flag-STUB1 in transfected HEK293T cells. (**G**) Mapping of the YTHDF1 domain binding to STUB1. Schematic depiction of domain-deleted YTHDF1 (upper). Co-IP assays of Myc-STUB1 and truncated Flag-YTHDF1 in transfected HEK293T cells.

We therefore proceeded to map the interaction domain of STUB1 and YTHDF1. STUB1 contains of three domains, including an N-terminal TPR domain, a central coiled-coil CC domain, and a U-box domain at C-terminal. Co-immunoprecipitation assays demonstrated that the TPR domain interacted with YTHDF1 while deletion of the TPR domain abolished STUB1 binding to YTHDF1, indicating the TPR domain of STUB1 mediated the interaction with YTHDF1 (Fig. 5F). Furthermore, we also constructed several domain-deleted Flag-YTHDF1 to determine the domain of YTHDF1 involved in STUB1 binding. Co-IP assays showed that YTH domain of YTHDF1 interacted with Myc-STUB1 (Fig. 5G). Taken together, these data demonstrate that the STUB1 TPR domain interact with the YTH domain of YTHDF1.

### STUB1 promotes ubiquitylation and degradation of YTHDF1

As STUB1 is an E3 ubiquitin ligase, we investigated whether STUB1 regulates YTHDF1 ubiquitination. As expected, knockdown of STUB1 inhibits, enforced expression of STUB1 promotes, YTHDF1 ubiquitylation in HEK293T cells (Fig. 6A, B). We next explored the ubiquitin chain types of YTHDF1 was regulated by STUB1. The results show that Lys 48-linked ubiquitylation of YTHDF1 was decreased by STUB1 knockdown, while the Lys 63-linked ubiquitylation displayed little change upon STUB1 knockdown (Fig. 6C, D).

**Figure 6.**
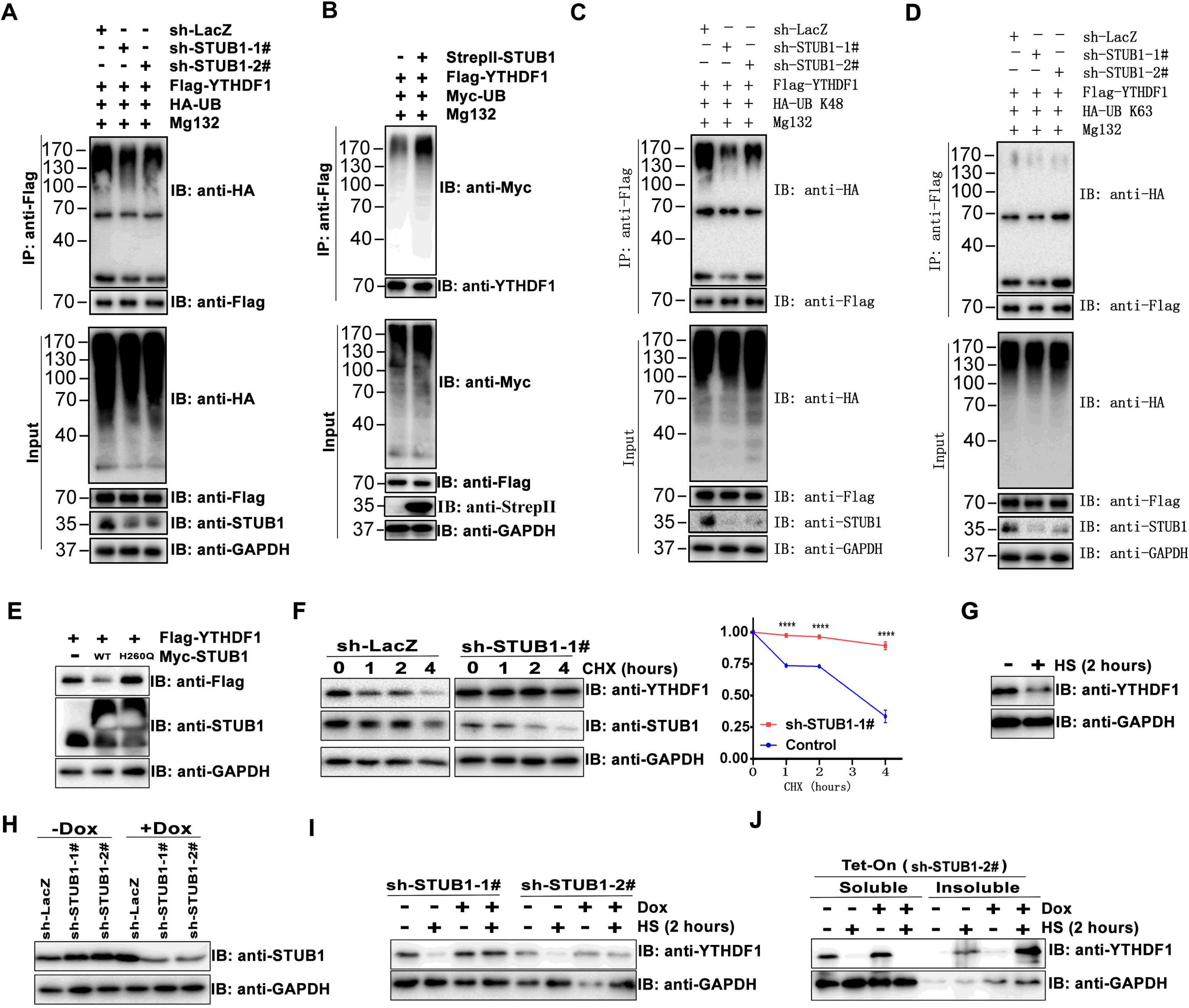
STUB1 promotes ubiquitylation and degradation of YTHDF1. (**A**) Knockdown of STUB1 inhibits YTHDF1 ubiquitylation in HEK293T cells. HEK293T were infected with both lentivirus expressing Flag-YTHDF1 and sh-LacZ/-STUB1-1#/-STUB1-2# to establish a stably expressing cell line, and then transfected with HA-UB. At 48 h after transfection, the cells were treated with MG132 (10 μg/mL) for 8 hours. Flag-YTHDF1 were pulled down by anti-Flag agarose under denaturing conditions, and then analyzed by Western blotting with HA antibodies. (**B**) enforced expression of STUB1 promotes YTHDF1 ubiquitylation in HEK293T cells. HEK293T cells were transfected with Flag-YTHDF1, Myc-UB, with or without StrepII-STUB1 as indicated. After 48 hours, the cells were treated with MG132 (10 μg/mL) for 8 hours. (**C, D**) HEK293T cells stably expressing Flag-YTHDF1 and sh-LacZ/-STUB1-1#/-STUB1-2# were transfected with HA tagged mutant Ub (K48, Lys 48-only; K63, Lys 63-only), and then treated with MG132 (10 μg/mL) for 8 hours. (**E**) HEK293T cells were co-transfected with Flag-YTHDF1 and Myc tagged WT STUB1 or mutant STUB1 (H260Q). Cells were lysed and immunoblotted using STUB1, Flag and GAPDH antibodies. (**F**) Cycloheximide (CHX) chase experiments showed that STUB1 knockdown prolongs YTHDF1 half-life. HEK293T cells stably expressing sh-LacZ or sh-STUB1 were treated with 100 μmol/L cycloheximide for the indicated time period. The YTHDF1 and STUB1 proteins were detected by immunoblots. Quantification of YTHDF1 protein levels shown on the right. (**G**) HEK293T were cultured under normal conditions or heat shock (HS) conditions for 2 hours, and then the cells were lysed and immunoblotted with YTHDF1. (**H**) Generation of doxycycline (dox)-inducible STUB1-knockdown HEK293T cells. The expression levels of STUB1 was determined by STUB1 antibody. GAPDH is an internal control. (**I**) Heat-shock mediated reduction of YTHDF1 protein level was block by STUB1 knockdown. Immunoblot for YTHDF1 in HEK293T cells stably expressing dox-inducible sh-STUB1-1# or sh-STUB1-2# after treatment with dox or/and HS as indicated. (**J**) HEK293T cells stably expressing dox-inducible sh-STUB1-2# was exposed to dox treatment for 72 hours, followed by heat shock treatment for 2 hours. The cells were lysed in NETN, and then separated into NETN-soluble and NETN-insoluble fractions, followed by western blotting with indicated antibodies.

We next sought to investigate whether STUB1 is involved in the regulation of YTHDF1 protein level. To so do, we co-transfected YTHDF1 with the full-length STUB1 (WT) or the H260Q mutant plasmid lacking ubiquitin ligase activity into HEK293T cells. We found overexpression of wide type STUB1 significantly reduced Flag-YTHDF1 protein levels, whereas STUB1-H260Q overexpression had little effect on Flag-YTHDF1 protein levels (Fig. 6E), indicating that STUB1 decrease YTHDF1 level requires its ubiquitin ligase activity. Consistently, cycloheximide (CHX) chase experiments showed that STUB1 knockdown prolongs YTHDF1 half-life (Fig. 6F). The above results indicate that STUB1 promotes the ubiquitination and degradation of YTHDF1.

Previous studies have shown that YTHDF1 exhibits dynamic protein aggregation and relocalization during heat shock^[31]^, and STUB1 plays an important role as a co-chaperone protein in the heat shock (HS) and aggregation pathways. We therefor investigated whether heat-shock could affect the protein levels of YTHDF1. As expected, the protein level of YTHDF1 significantly decreased when the cells were treated with heat shock for 2 h (Fig. 6G). We also asked whether STUB1 knockdown affects heat-shock induced decrease of YTHDF1 protein level. To this end, we generated HEK293T cells stably expressing doxycycline (dox)-inducible shRNAs targeting STUB1 and controls targeting LacZ. Dox treatment successful induced the down-regulation of STUB1 protein level (Fig. 6H). In addition, we found that heat-shock mediated reduction of YTHDF1 was block by STUB1 knockdown (Fig. 6I). Furthermore, we found that heat-shock stress led to the accumulation of YTHDF1 in the insoluble aggresome (Fig. 6J); and STUB1 knockdown resulted in further YTHDF1 accumulation in the insoluble fraction compared with that in the control cells (Fig. 6J). Thus, heat shock proteins may contribute to STUB1-mediated YTHDF1 degradation.

### STUB1 inhibits renal cancer cell migration and invasion though degrading YTHDF1

To explore the possible function of YTHDF1 in ccRCC tumorigenesis, we knocked down YTHDF1 level with two distinct dox-inducible shRNAs in 786-O and OS-RC-2 cells (Fig. 7A), and examined the effect of YTHDF1 knockdown on cell proliferation. The result showed that two independent YTHDF1 shRNAs significantly decreased proliferation of these two cells (Fig. 7B). Moreover, the transwell migration/invasion assays showed that the migration and invasion capability of 786-O and OS-RC2 cells were significantly reduced after knocking down YTHDF1 (Fig. 7C-E). Thus, YTHDF1 functions as an oncogene to promote cell growth, migration, and invasion of renal cancer cells.

**Figure 7.**
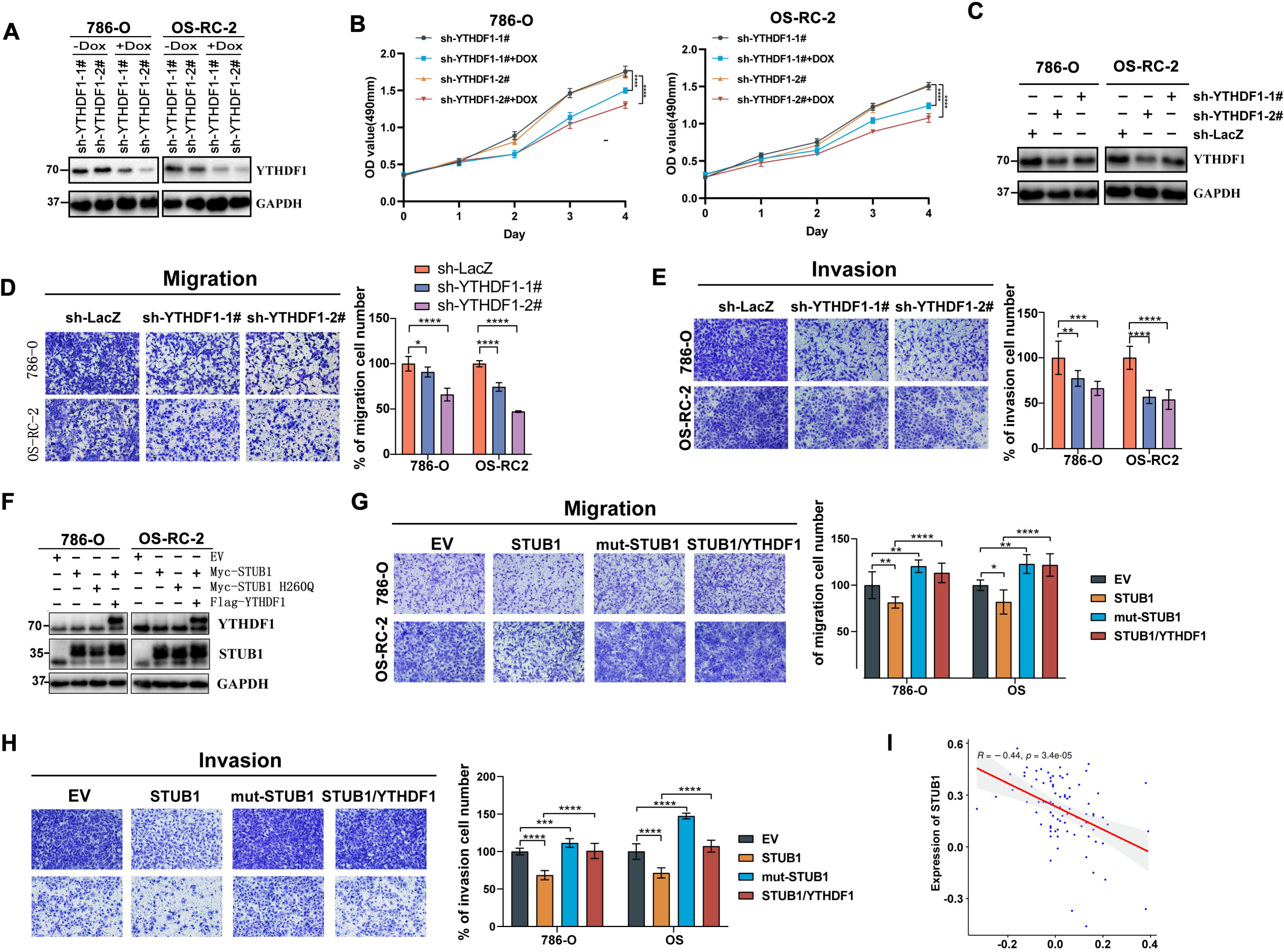
STUB1 inhibits renal cancer cell migration and invasion though degrading YTHDF1. (**A**) Generation of doxycycline (dox)-inducible YTHDF1-knockdown 786-O and OS-RC-2 cells. The expression level of YTHDF1 was evaluated by western blotting. (**B**) MTS assay for 786-O and OS-RC-2 cell lines with stable expression of dox-inducible sh-YTHDF1-1# or sh-YTHDF1-2#. (**C**) Generation of stably YTHDF1-knockdown 786-O and OS-RC-2 cells. The expression level of YTHDF1 was evaluated by western blotting. (**D, E**) Migration (D) and invasion (E) assays for 786-O and OS-RC-2 cell lines with stable expression sh-LacZ, sh-YTHDF1-1# or sh-YTHDF1-2#. Migrated and invaded cells from each treatment group were counted in four random images. (**F**) 786-O and OS-RC-2 cells were transfected with Myc-STUB1 or Myc-STUB1 H260Q along with Flag-YTHDF1 as indicated. The YTHDF1 and STUB1 proteins were detected by immunoblots. (**G, H**) Migration (G) and invasion (H) assays for 786-O and OS-RC-2 cell lines stably expressing plasmids as in panel F, cells were then used for transwell assays. (**I**) STUB1 protein showed a significantly negative correlation with YTHDF1 protein in normal adjacent tissues. Based on the CPTAC dataset, we analyzed the expression level of STUB1 and YTHDF1 in normal adjacent tissues of ccRCC tumor. EV: Empty vector.

Next, we tested whether YTHDF1 is required for STUB1-mediated cancer-inhibitory effects in renal cancer cells. We found that forced expression of YTHDF1 restored the migration and invasion of STUB1-overexpression cells (Fig. 7F-H), suggesting that STUB1 inhibits migration and invasion in renal cancer cell is mediated, at least in part, through ubiquitinating YTHDF1.

We further performed an analysis of ccRCC datasets from the CPTAC database and found that STUB1 protein showed a significantly negative correlation with YTHDF1 protein in normal adjacent tissues (Figure 7I). However, the STUB1 protein levels in tumor tissues showed no correlation between YTHDF1 protein levels (data not shown), suggesting that other regulators may also regulate the YTHDF1 expression in the ccRCC tumor tissues.

## Discussion

This study provides evidence that STUB1 is frequently down-regulated in ccRCC and functions as a tumor suppressor. Mechanistically, we demonstrated that STUB1 can promote the ubiquitination and degradation of YTHDF1. Importantly, our results showed that STUB1 inhibits migration and invasion in renal cancer cell is mediated, at least in part, through ubiquitinating YTHDF1.

Previous studies have demonstrated that the E3 ubiquitin ligase STUB1 exhibits a tumor suppressor effect by promoting ubiquitination and degradation of multiple oncogenes in various cancers, including lung cancer^[16, 17]^, breast cancer^[7]^, prostate cancer^[8-10]^, gastric cancer^[21]^, colorectal cancer^[14]^, and thyroid cancer^[32]^. For example, STUB1 functions as a tumor suppressor in ovarian carcinoma by promoting ubiquitination and degradation of PKM2^[19]^. In addition, it has also been shown that STUB1 can ubiquitinate and degrade specific tumor suppressors, thus functions as an oncogene in certain tumors^[24-26, 33]^. For example, STUB1 can promote the activations of AKT, NF-kB, and WNT signaling pathways in breast cancer cells by ubiquitinating PFN1, therapy promotes breast cancer metastasis^[24]^. Thus, STUB1 is an important E3 ubiquitin ligase in regulating tumor development and progression. Sun *et al*. showed that STUB1 expression is down-regulated in ccRCC tissues, and its low expression is associated with poor survival of ccRCC patients^[34]^. Min *et al*. demonstrated that STUB1 suppress renal cancer cell growth and angiogenesis by targeting TG2 for degradation^[20]^. In this study, we found that STUB1 interacted with YTHDF1 and enhanced its ubiquitination and degradation, which reinforced that STUB1 play an important role in ccRCC. Therefore, our results point that STUB1 may be a therapeutic target in ccRCC.

STUB1 consists of three tetratricopeptide-repeat (TPR) domains at N-terminal that mediate the interaction with chaperone, a central coiled-coil CC domain and a C-terminal U-box domain that recruits ubiquitin-conjugating enzymes (E2s). STUB1 is able to link molecular chaperones to the ubiquitin-proteasome system (UPS), thereby playing a key role in protein quality control^[35]^. It has been demonstrated that STUB1 recognizes the substrates in both molecular chaperones-dependent and -independent manner. For example, STUB1 interacts with OTUD3 though its CC domain, and targets the latter for degradation^[16]^. STUB1 interacts directly with SirT6 though the U-box domain and ubiquitinates SirT6^[36]^. Here we screened interaction proteins of STUB1 using AP-MS, and identified YTHDF1 as a new STUB1 binding partner. We further confirmed the interaction between STUB1 and YTHDF1 by co-immunoprecipitation and GST pull down assays, suggesting STUB1 interacts directly with YTHDF1 and is not dependent on heat shock proteins (HSPs). In addition, we also found heat shock stress induced the degradation of YTHDF1, and demonstrated that STUB1 is required for the down-regulation of YTHDF1 upon heat shock stress. These results indicated that STUB1 promotes YTHDF1 ubiquitylation and degradation in HSPs dependent manner. It is possible that HSPs can promote the interaction of STUB1 with YTHDF1. Future work will focus on identifying which heat shock proteins involved in STUB1-mediated ubiquitylation of YTHDF1.

During our manuscript preparation, Liao *et al*. reported that YTHDF2 can interact with STUB1 and HSP90β^[37]^. In addition, their results showed that STUB1 induces the ubiquitination of YTHDF2, whereas HSP90β blocked the STUB1-mediated ubiquitination. Moreover, they revealed that the N-terminal (1-384 aa) of YTHDF2 is required for its binding to HSP90β. In our experiments, we found that the YTH domain of YTHDF1 is sufficient for STUB1 binding. Considering that the sequence and the structure of YTH domain of YTHDF1, YTHDF2 and YTHDF3 are highly conserved, it possible that all three YTHDF proteins can interact with STUB1 though their YTH domain. However, we found that the three YTHDF proteins have different abilities to bind STUB1, with YTHDF1 having the strongest binding ability to STUB1. The simplest explanation for these observations is that other regulators exist in modulating the interaction between YTHDF proteins and STUB1.

As a crucial m6A reader protein, YTHDF1 plays a pivotal role in various biology processes in tumor regulation, such as tumor immunity, drug resistance, tumor cell metabolism, proliferation, invasion, and metastasis^[38]^. Numerous studies have demonstrated that YTHDF1 expression is elevated in multiple cancers, and its overexpression is indicative of poor prognosis in tumor patients, thus highlighting its tumor-promoting role^[39-42]^. In this study, we provide functional evidence that YTHDF1 promotes RCC cell proliferation, migration and invasion *in vitro*. Although a number of studies have shown that YTHDF1 has a key regulatory role in tumors, its underlying molecular mechanisms have not been fully elucidated, and there exist some contrary findings. Studies have shown that YTHDF1 have different roles in different tumors, and even in the same tumor, different studies have reached opposite conclusions. For example, Tian *et al*. demonstrated that YTHDF1 could increase the protein level of tumor suppressor ANKLE1 in colorectal cancer (CRC) though m6A methylation^[43]^. However, YTHDF1 can also promote CRC development and metastasis by enhancing the translation of ARHGEF2, TCF7L2, and GLS1^[39, 44, 45]^. It is possible that YTHDF1’s opposite effect on tumor is caused by tumor heterogeneity; the same tumor exhibits many different genotypes or subtypes of cancer cells. Therefore, it will be important to determine the context-dependent functions of YTHDF1.

Post-translational modification (PTM) plays a key role in moderating YTHDF family proteins. For example, O-linked GlcNAc (O-GlcNAc) at S197 stimulate the cytosolic localization of YTHDF1^[46]^. The data presented here supports the conclusion that STUB1 is an E3 ubiquitin ligase of YTHDF1. Firstly, we confirmed the directly interaction between STUB1 and YTHDF1 through co-immunoprecipitation analysis and GST pull down assay. Secondly, STUB1 negatively regulates the protein level of YTHDF1 and is dependent on its ubiquitin ligase activity. Thirdly, STUB1 depletion inhibits the Lys48 linked polyubiquitination of YTHDF1. Although the direct association between STUB1 and YTHDF1 would indicate a direct mechanism for STUB1’s effect on the ubiquitylation of YTHDF1. However, it is possible that STUB1 promotes the ubiquitylation of YTHDF1 through an indirect mechanism, such as promoting the degradation of another E3 ligase that ubiquitinates YTHDF1. YTHDF1 harbors 16 lysine residues, including 3 lysines within the YTH domain. These lysine residues not only represent potential sites for ubiquitination, but also potential for many other post-translational modifications, including sumoylation and acetylation. Currently, the STUB1-catalyzed ubiquitination sites on YTHDF1 are still unknown. Therefore, future work will focus on identifying the ubiquitination sites on YTHDF1 using mass spectrometry methods, and exploring other regulators involved in YTHDF1 stability and their potential regulation by STUB1.

In summary, our study proposes a mechanism by which the tumor suppressor STUB1 interacts with YTHDF1 in renal cancer cells. Downregulation of STUB1 may result in the stability and abnormal upregulation of YTHDF1, which in turn contributes to the tumorigenic effects in RCC. These findings hold significant implications for the diagnosis and treatment of RCC.

## Materials and methods

### Human samples

The cancerous and paracancerous tissues used in this study were obtained from patients who underwent surgical treatment at Tongji Hospital (Wuhan, Hubei, China). Prior to the study, informed consent was obtained from all patients.

### Plasmids

The Myc-UB, HA-UB, HA-UB K48, or HA-UB K63 plasmids have been described previously^[47]^. The human YTHDF1, YTHDF2 and YTHDF3 cloned into pcDNA3.1-Flag-HA were from Prof. Shuliang Chen (Wuhan University, China). To generate StrepII-STUB1 and Myc-STUB1, the 5’ and 3’ primers of STUB1 were amplified by PCR to obtain STUB1 CDS, and then the amplified STUB1 was ligated into the pHAGE-StrepII and pHAGE-3xMyc plasmids. The CDS of STUB1 was amplified and cloned into psi-Cherry to generate STUB1-Cherry plasmid. Flag-HA-YTHDF1 was PCR amplified using primers YTHDF1-5’ and YTHDF1-3’, digested by BamHI and XhoI, and ligated into psi-Flag-C1, psi-HA, pEGFP-myc-C1, pHAGE-3xMyc, pGEX-4T-1 to create Flag-YTHDF1, HA-YTHDF1, EGFP-YTHDF1, Myc-YTHDF1 lentiviral vectors and GST-YTHDF1, respectively. The STUB1 H260Q point mutant was generated by a two-step PCR-based mutagenesis procedure using Myc-STUB1 plasmid as the template. The truncated mutants of STUB1 and YTHDF1 were constructed by standard molecular biology techniques. The primers used for PCR amplification are listed in Supplementary Table S2.

Lenti-H1TO, is a dox-inducible shRNA expression vector, was constructed by inserting the minimal H1/TO promoter, the EF-1α core promoter and the tetracycline repressor (TetR) into the KpnI and BamHI sites of lentiCRISPR-v2 (Addgene, 52961). Then, shRNA sequences corresponding to the target sequences were ligated into the Age1 and NheI sites of the Lenti-H1/TO plasmid. The following target regions were chosen: STUB1-1#: 5’-GGCCTTGTGCTACCTGAAGAT-3’; STUB1-2#: 5’-GAAGCGCTGGAACAGCATTGA-3’; YTHDF1-1#: 5’-GCACTGACTGGTGTCCTTTCT-3’; YTHDF1-2#: 5’-GGGTCTGGTCTCAGGACAAGT-3’.

### Antibodies

The antibodies employed in this experiment were as follows: mouse anti-glyceraldehyde-3-phosphate dehydrogenase (GAPDH) (G8795, 1:4000), anti-Flag (F3165, 1:2000) from Sigma-Aldrich (St. Louis, MO, USA); mouse anti-hemagglutinin (HA) antibody (11583816001, 1:3000) from Roche Applied Science (Penzberg, Germany); rabbit anti-Myc-tag (562, 1:3000) and mouse anti-Strep-tag II (M211-3, 1:3000) from MBL Life Science (MBL, Nagoya, Japan); rabbit anti-STUB1 (A11751, 1:3000) from ABclonal (Wuhan, China); rabbit anti-YTHDF1 (17479-1-AP, 1:3000) from Proteintech (Wuhan, China); goat anti-mouse IgG-horseradish peroxidase (HRP) secondary antibody (31430, 1:20,000); and goat anti-rabbit IgG-horseradish peroxidase (HRP) secondary antibody (31460, 1:20,000) from Thermo Fisher Scientific (Waltham, MA, USA).

### Cell culture

The ccRCC cell lines 786-O and OS-RC-2 cells were cultured in RPMI-1640 medium (White Shark Biologicals, Anhui, China) supplemented with 10% fetal bovine serum (Gibco, South America). The human embryonic kidney cells HEK293T cells were cultured in DMEM medium (White Shark Biologicals, Anhui, China) supplemented with 5% fetal bovine serum. Cell cultures were regularly shown to be free of mycoplasma.

### Lentivirus packaging and infection, cell transfections

To achieve lentiviral transfection, HEK293T cells were co-transfected with packaging plasmids PDM2G and PSPAX2, as well as sh-LacZ, shRNA STUB1, shRNA YTHDF1, or overexpression plasmids for STUB1 and YTHDF1. After 48 hours, the supernatant containing the virus was collected and used to infect 786-O and OS-RC-2 cells, as well as HEK293T cells. The stably transfected cell lines were then screened with 2 μg/ml puromycin or 300 μg/ml hygromycin.

### Immunohistochemistry (IHC)

Formalin-fixed and paraffin-embedded surgical specimens were sectioned and deparaffinized using a xylene solution and a gradient of ethanol solutions. The sections were then washed with 1× Tris-buffered saline (TBS) or 1× phosphate-buffered saline (PBS). Antigen retrieval was performed by heating the sections in a microwave oven for 10 minutes with 0.1M sodium citrate. Non-specific binding was blocked with A blocking reagent (Gene Technology) (GK600505, Shanghai, China). The primary antibody used was rabbit anti-STUB1 (A11751, 1:3000), and the secondary antibody was Goat Anti-Mouse/Anti-Rabbit (Gene Technology). The chromogenic agent used was also obtained from Gene Technology.

### RT-PCR and RT-qPCR

Total RNA was extracted using Trizol reagent (Vazyme, China), and cDNA was synthesized using a highly efficient second-generation RT-qPCR reagent (Vazyme, China) in premix form. Real-time fluorescence quantitative PCR was performed using the SYBR Mix, following standard protocols. The primers for RT-qPCR are shown in Supplementary Table S2.

### Co-immunoprecipatation (Co-IP) and GST-pull down

Protein interactions were analyzed using the Co-IP assays. Cells were transfected with indicated plasmids and lysed in NETN buffer. Appropriate protein antibodies were added to the cell lysate, which was then gently rotated and mixed for 1-2 hours at 4°C. After that, Protein A/G agarose beads were added, and the mixture was incubated at 4°C overnight. The samples were washed 3-5 times with NETN buffer supplement with 150 mM NaCl, and the bound proteins were eluted using Elution Buffer. Subsequently, they were separated by sodium dodecyl sulfate polyacrylamide gel electrophoresis (SDS-PAGE).

E. coli BL21 cells containing GST-YTHDF1 or pGEX-4T-1 plasmid were cultured and induced with IPTG for 16 hours, and then the GST-YTHDF1 and GST were purified with glutathione beads (GE Healthcare). The resulting purified GST-YTHDF1 or GST was incubated with Myc-STUB1 purified from HEK293T cells. Then, the bead fraction was extensive washed and analyzed by western blotting.

### AP-MS

Approximately 100 million cells stably expressing StrepII or StrepII-STUB1 were lysed in NETN lysis buffer. The extract was incubated with Strep-Tactin XT superfluid resin (2-4010; IBA Lifesciences, Gottingen, Germany) at 4°C for 4 hours. Then, the beads were washed five times with wash buffer and eluted with NuPAGE™ LDS sample buffer (NP 0007; Thermo Fisher Scientific) followed by SDS-PAGE and Coomassie blue staining. In-gel proteins were digested and analyzed using an HPLC-Orbitrap-Elite mass spectrometer (Thermo Fisher Scientific).

### Cycloheximide chase assays

Cells were cultured in 12-well plates and when they reached 80%-90% confluency, they were treated with cycloheximide (CHX, A10036, AdooQ BioScience, Irvine, CA, USA) at a final concentration of 100 μmol/L. The cells were then harvested at indicated time points after CHX treatment and subjected to immunoblotting with specific or control antibodies.

### Ubiquitination assays

HEK293T cells stably transfected with Flag-YTHDF1 were further stably transfected with sh-LacZ, sh-STUB1-1#, and sh-STUB1-2#. The resulting cells were then transiently transfected with HA-UB, HA-UB K48, or HA-UB K63. Approximately 48 h after transfection, MG 132 was added at a final concentration of 10 μg/mL for 8 hours prior to cell lysis using 3% SDS Lysis Buffer. The lysed cells were boiled in water for 10 minutes and then incubated with NETN lysis buffer at 4°C for 1 hour. Then, the supernatant was incubated with a Flag antibody and Protein A/G Agarose for Co-IP experiments. The bound proteins were then subjected to western blot analysis.

### Cell proliferation, migration and invasion assay

For clone formation experiments, 1500 cells were grown in each well of a 6-well plate. After 11-14 days of growth, the cells were fixed with methanol for half an hour. Then, the cells were stained with a 0.5% crystal violet solution for half an hour and washed with water. The colonies in the 6-well plates were photographed and counted. In the MTS experiment, 96-well plates were filled with approximately 2000 cells per well and incubated with the MTS reagent for a specified duration as per the instructions. The OD value was measured at a wavelength of 490nm using a microplate reader, while being incubated in the dark. In migration and invasion assays, 30,000-50,000 cells are suspended in 250ul of serum-free medium and added to Transwell chambers. The inserts were then placed in wells of a 24-well plate containing 500 μL serum-containing medium. After 12-24 hours, the cells were fixed with methanol for 30 minutes and stained with 5% crystal violet for 45 minutes.

### Animal experiments

A total of 2×10^6^ cells suspended in 100 μL PBS solution were subcutaneously injected into about 7-week-old male BALB/C nude mice obtained from Beijing Huafukang Biotechnology (Beijing, China). One week after the injection (day 8), the mice were placed under Doxycycline (Dox) liquid diet (1 mg/ml Dox and 5% sucrose), and the water was replaced every 3 days. The length, width, and height of the tumor were measured using a caliper, and tumor volume was calculated using the formula: 0.52×length×width×height. Finally, the tumor was weighed using an electronic scale. Animal experiments were approved by the Animal Ethics Committee of Wuhan Tongji Hospital.

### TCGA and CPTAC analysis

Transcriptome and clinical data related to kidney cancer were obtained from the TCGA database (https://portal.gdc.cancer.gov/). R packages including “ggsci”, “survival”, and “survminer” were utilized to group the data based on clinical characteristics and create boxplots. The prognosis of patients was analyzed according to the STUB1 expression level.

The complete proteomic TMT data of ccRCC patients was obtained from the CPTAC database (https://pdc.cancer.gov/pdc/). Paired sample analysis was performed using the R package “ggpubr”, and paired sample boxplots were constructed.

## Supporting information

Supplementary Figure S1. Immunohistochemical results of tumor tissues of 12 pairs of kidney cancer patients and their paired paracancerous tissues.

Supplementary Table S1. Proteins found in the StrepII-STUB1 precipitates according to LC-MS analysis

Supplementary Table S2. The primers used in the study

## Statistical analysis

Comparison of the means of the two groups was performed using Student’s t-test. For comparison between the means of multiple groups, a one-way ANOVA was selected. The Kaplan-Meier curves and the log-rank test was used for comparing the survival times obtained from two groups.

## Author contributions

Z.Z., K.C. conceived and planned experiments, S.M. and K.C. performed the AP-MS screening, Co-IP, and Western blot analyses, Y.S. and J.Z. did the cell culturing and qPCR analyses. G.G. and S.M. performed data analysis and the statistical analysis. S.M., Z.Z., and K.C. contributed to manuscript writing. All authors edited and approved the manuscript.

## Acknowledgements

Not applicable.

## Ethics approval and consent to participate

This study was approved by the Tongji hospital of Tongji Medical College, Huazhong University of Science and Technology (Wuhan, China) ethics review committee. All human tumor tissues were obtained with written informed consent from patients prior to participation in the study.

## Data Availability

The dataset generated and/or analyzed during the current study is not publicly available but may be available from the corresponding author upon reasonable request.

## References

1. Sato, Y., T. Yoshizato, Y. Shiraishi, et al., Integrated molecular analysis of clear-cell renal cell carcinoma. Nat Genet, 2013. 45(8): p. 860–7.

2. Rosellini, M., A. Marchetti, V. Mollica, et al., Prognostic and predictive biomarkers for immunotherapy in advanced renal cell carcinoma. Nat Rev Urol, 2023. 20(3): p. 133–157.

3. Braun, D.A., Z. Bakouny, L. Hirsch, et al., Beyond conventional immune-checkpoint inhibition - novel immunotherapies for renal cell carcinoma. Nat Rev Clin Oncol, 2021. 18(4): p. 199–214.

4. Choueiri, T.K. and R.J. Motzer, Systemic Therapy for Metastatic Renal-Cell Carcinoma. N Engl J Med, 2017. 376(4): p. 354–366.

5. Kumar, S., M. Basu, and M.K. Ghosh, Chaperone-assisted E3 ligase CHIP: A double agent in cancer. Genes Dis, 2022. 9(6): p. 1521–1555.

6. Kajiro, M., R. Hirota, Y. Nakajima, et al., The ubiquitin ligase CHIP acts as an upstream regulator of oncogenic pathways. Nat Cell Biol, 2009. 11(3): p. 312–9.

7. Luan, H., B. Mohapatra, T.A. Bielecki, et al., Loss of the Nuclear Pool of Ubiquitin Ligase CHIP/STUB1 in Breast Cancer Unleashes the MZF1-Cathepsin Pro-oncogenic Program. Cancer Res, 2018. 78(10): p. 2524–2535.

8. Xu, S., L. Fan, H.Y. Jeon, et al., p300-Mediated Acetylation of Histone Demethylase JMJD1A Prevents Its Degradation by Ubiquitin Ligase STUB1 and Enhances Its Activity in Prostate Cancer. Cancer Res, 2020. 80(15): p. 3074–3087.

9. Liu, C., W. Lou, J.C. Yang, et al., Proteostasis by STUB1/HSP70 complex controls sensitivity to androgen receptor targeted therapy in advanced prostate cancer. Nat Commun, 2018. 9(1): p. 4700.

10. Sarkar, S., D.L. Brautigan, S.J. Parsons, et al., Androgen receptor degradation by the E3 ligase CHIP modulates mitotic arrest in prostate cancer cells. Oncogene, 2014. 33(1): p. 26–33.

11. Sarkar, S., D.L. Brautigan, and J.M. Larner, Aurora Kinase A Promotes AR Degradation via the E3 Ligase CHIP. Mol Cancer Res, 2017. 15(8): p. 1063–1072.

12. Wang, J., H. Zhang, X. Zhang, et al., PC-1 works in conjunction with E3 ligase CHIP to regulate androgen receptor stability and activity. Oncotarget, 2016. 7(49): p. 81377–81388.

13. Zhong, Y., T. Long, C.S. Gu, et al., MYH9-dependent polarization of ATG9B promotes colorectal cancer metastasis by accelerating focal adhesion assembly. Cell Death Differ, 2021. 28(12): p. 3251–3269.

14. Wang, Y., F. Ren, Y. Wang, et al., CHIP/Stub1 functions as a tumor suppressor and represses NF-κB-mediated signaling in colorectal cancer. Carcinogenesis, 2014. 35(5): p. 983–91.

15. Wang, J., Y. Zhou, D. Zhang, et al., CRIP1 suppresses BBOX1-mediated carnitine metabolism to promote stemness in hepatocellular carcinoma. Embo j, 2022. 41(15): p. e110218.

16. Zhang, P., C. Li, H. Li, et al., Ubiquitin ligase CHIP regulates OTUD3 stability and suppresses tumour metastasis in lung cancer. Cell Death Differ, 2020. 27(11): p. 3177–3195.

17. Shi, Y., X. Wang, Z. Xu, et al., PDLIM5 inhibits STUB1-mediated degradation of SMAD3 and promotes the migration and invasion of lung cancer cells. J Biol Chem, 2020. 295(40): p. 13798–13811.

18. Wang, T., J. Yang, J. Xu, et al., CHIP is a novel tumor suppressor in pancreatic cancer through targeting EGFR. Oncotarget, 2014. 5(7): p. 1969–86.

19. Shang, Y., J. He, Y. Wang, et al., CHIP/Stub1 regulates the Warburg effect by promoting degradation of PKM2 in ovarian carcinoma. Oncogene, 2017. 36(29): p. 4191–4200.

20. Min, B., H. Park, S. Lee, et al., CHIP-mediated degradation of transglutaminase 2 negatively regulates tumor growth and angiogenesis in renal cancer. Oncogene, 2016. 35(28): p. 3718–28.

21. Tang, D.E., Y. Dai, L.W. Lin, et al., STUB1 suppresseses tumorigenesis and chemoresistance through antagonizing YAP1 signaling. Cancer Sci, 2019. 110(10): p. 3145–3156.

22. Ahmed, S.F., N. Das, M. Sarkar, et al., Exosome-mediated delivery of the intrinsic C-terminus domain of PTEN protects it from proteasomal degradation and ablates tumorigenesis. Mol Ther, 2015. 23(2): p. 255–69.

23. Choi, J.R., K.S. Shin, C.Y. Choi, et al., PARP1 regulates the protein stability and proapoptotic function of HIPK2. Cell Death Dis, 2016. 7(10): p. e2438.

24. Zhang, S., X. Guo, X. Liu, et al., Adaptor SH3BGRL promotes breast cancer metastasis through PFN1 degradation by translational STUB1 upregulation. Oncogene, 2021. 40(38): p. 5677–5690.

25. Han, S.Y., A. Ko, H. Kitano, et al., Molecular Chaperone HSP90 Is Necessary to Prevent Cellular Senescence via Lysosomal Degradation of p14ARF. Cancer Res, 2017. 77(2): p. 343–354.

26. Li, F., P. Xie, Y. Fan, et al., C terminus of Hsc70-interacting protein promotes smooth muscle cell proliferation and survival through ubiquitin-mediated degradation of FoxO1. J Biol Chem, 2009. 284(30): p. 20090–8.

27. Biswas, K., S. Sarkar, K. Du, et al., The E3 Ligase CHIP Mediates p21 Degradation to Maintain Radioresistance. Mol Cancer Res, 2017. 15(6): p. 651–659.

28. Liang, Z.L., M. Kim, S.M. Huang, et al., Expression of carboxyl terminus of Hsp70-interacting protein (CHIP) indicates poor prognosis in human gallbladder carcinoma. Oncol Lett, 2013. 5(3): p. 813–818.

29. Clark, D.J., S.M. Dhanasekaran, F. Petralia, et al., Integrated Proteogenomic Characterization of Clear Cell Renal Cell Carcinoma. Cell, 2019. 179(4): p. 964-983.e31.

30. Qu, Y., J. Feng, X. Wu, et al., A proteogenomic analysis of clear cell renal cell carcinoma in a Chinese population. Nat Commun, 2022. 13(1): p. 2052.

31. Ries, R.J., S. Zaccara, P. Klein, et al., m(6)A enhances the phase separation potential of mRNA. Nature, 2019. 571(7765): p. 424–428.

32. Xu, Y., G. Xu, H. Dang, et al., Carboxy terminus of HSP70-interacting protein (CHIP) attenuates the stemness of thyroid cancer cells through decreasing OCT4 protein stability. Environ Toxicol, 2021. 36(4): p. 686–693.

33. Tanwar, K. and U. Pati, Inhibition of apoptosis via CHIP-mediated proteasomal degradation of TAp73α. J Cell Biochem, 2019. 120(7): p. 11091–11103.

34. Sun, C., H.L. Li, H.R. Chen, et al., Decreased expression of CHIP leads to increased angiogenesis via VEGF-VEGFR2 pathway and poor prognosis in human renal cell carcinoma. Sci Rep, 2015. 5: p. 9774.

35. Liu, Y., H. Zhou, and X. Tang, STUB1/CHIP: New insights in cancer and immunity. Biomed Pharmacother, 2023. 165: p. 115190.

36. Ronnebaum, S.M., Y. Wu, H. McDonough, et al., The ubiquitin ligase CHIP prevents SirT6 degradation through noncanonical ubiquitination. Mol Cell Biol, 2013. 33(22): p. 4461–72.

37. Liao, Y., Y. Liu, C. Yu, et al., HSP90β Impedes STUB1-Induced Ubiquitination of YTHDF2 to Drive Sorafenib Resistance in Hepatocellular Carcinoma. Adv Sci (Weinh), 2023: p. e2302025.

38. Chen, Z., X. Zhong, M. Xia, et al., The roles and mechanisms of the m6A reader protein YTHDF1 in tumor biology and human diseases. Mol Ther Nucleic Acids, 2021. 26: p. 1270–1279.

39. Wang, S., S. Gao, Y. Zeng, et al., N6-Methyladenosine Reader YTHDF1 Promotes ARHGEF2 Translation and RhoA Signaling in Colorectal Cancer. Gastroenterology, 2022. 162(4): p. 1183–1196.

40. Shi, Y., S. Fan, M. Wu, et al., YTHDF1 links hypoxia adaptation and non-small cell lung cancer progression. Nat Commun, 2019. 10(1): p. 4892.

41. Sun, Y., D. Dong, Y. Xia, et al., YTHDF1 promotes breast cancer cell growth, DNA damage repair and chemoresistance. Cell Death Dis, 2022. 13(3): p. 230.

42. Li, Q., Y. Ni, L. Zhang, et al., HIF-1α-induced expression of m6A reader YTHDF1 drives hypoxia-induced autophagy and malignancy of hepatocellular carcinoma by promoting ATG2A and ATG14 translation. Signal Transduct Target Ther, 2021. 6(1): p. 76.

43. Tian, J., P. Ying, J. Ke, et al., ANKLE1 N(6) -Methyladenosine-related variant is associated with colorectal cancer risk by maintaining the genomic stability. Int J Cancer, 2020. 146(12): p. 3281–3293.

44. Han, B., S. Yan, S. Wei, et al., YTHDF1-mediated translation amplifies Wnt-driven intestinal stemness. EMBO Rep, 2020. 21(4): p. e49229.

45. Chen, P., X.Q. Liu, X. Lin, et al., Targeting YTHDF1 effectively re-sensitizes cisplatin-resistant colon cancer cells by modulating GLS-mediated glutamine metabolism. Mol Ther Oncolytics, 2021. 20: p. 228–239.

46. Li, J., M. Ahmad, L. Sang, et al., O-GlcNAcylation promotes the cytosolic localization of the m(6)A reader YTHDF1 and colorectal cancer tumorigenesis. J Biol Chem, 2023. 299(6): p. 104738.

47. Chen, K., J. Zeng, H. Xiao, et al., Regulation of glucose metabolism by p62/SQSTM1 through HIF1α. J Cell Sci, 2016. 129(4): p. 817–30.

