## Supplementary Figure S1. Immunohistochemical results of tumor tissues of 12 pairs of kidney cancer patients and their paired paracancerous tissues. for "The ubiquitin ligase STUB1 suppresses tumorigenesis of renal cell carcinomas through regulating YTHDF1 stability"

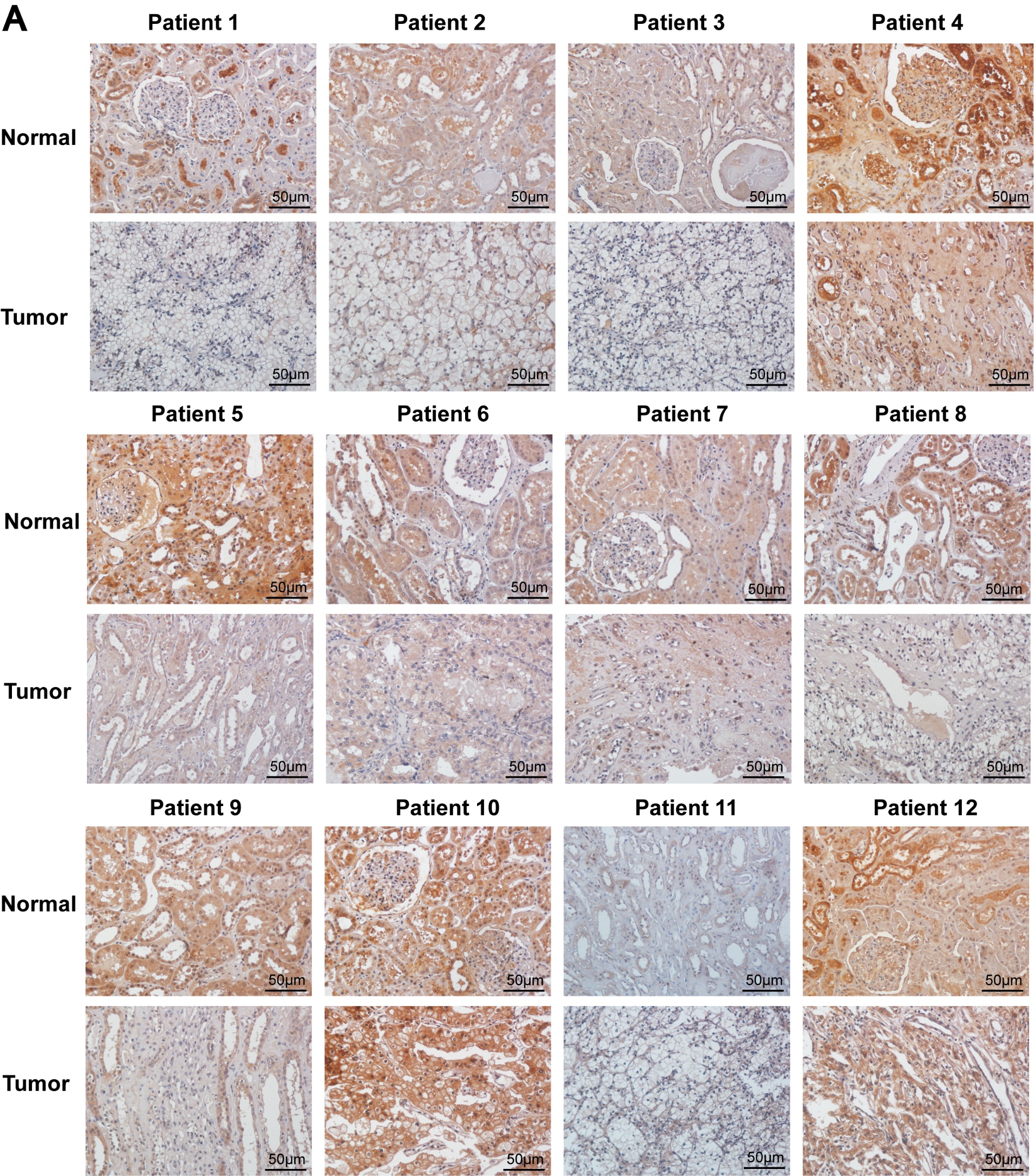

Figure S1. Immunohistochemical results of tumor tissues of 12 pairs of kidney cancer patients and their paired paracancerous tissues.
