## Supplementary Table S2. The primers used in the study for "The ubiquitin ligase STUB1 suppresses tumorigenesis of renal cell carcinomas through regulating YTHDF1 stability"

Sites for restriction enzymes are underlined.

| **Primer name** | **Primer sequence (5′-3′)** | **PCR** |
| --- | --- | --- |
| **Primers for making constructs** | | |
| STUB1-5'  STUB1-3' | CGCGGATCCATGAAGGGCAAGGAGGAGA  CCGCTCGAGGTAGTCCTCCACCCAGCCA | 94 °C, 30 s; 58 °C, 30 s; 72 °C, 120 s; 35 cycles |
| YTHDF1-5'  YTHDF1-3' | CGCGGATCCATGTCGGCCACCAGCGTGGA  ACGCGTCGACTTGTTTGTTTCGACTCTGCCG | 94 °C, 30 s; 58 °C, 30 s; 72 °C, 120 s; 35 cycles |
| STUB1(1-127)-3'  STUB1(225-330)-5'  STUB1(125-303)-5' | CCGCTCGAGCTGCTCCTTGGCCAGGCTGTA  CGCGGATCCCGAGACATCCCCGACTACCT  CGCGGATCCAAGGAGCAGCGGCTGAACT | 94 °C, 30 s; 58 °C, 30 s; 72 °C, 30 s; 35 cycles |
| YTHDF1(1-124) -3'  YTHDF1(1-365) -3'  YTHDF1(124-193) -5'  YTHDF1(124-193) -3'  YTHDF1(193-365) -5'  YTHDF1(365-559) -5'  YTHDF1(365-523) -3' | CCGCTCGAGAGGGAAAAAATTGAACCTGTG  ACGCGTCGACGTGGGATTCGACGCTGGG  CGCGGATCCCCTGAAAACCCTGCGTTCT  CCGCTCGAGCCCAATCTTCAGGCCAACC  CGCGGATCCGGGGACGTCAGCTCCTC  CGCGGATCCCACCCCGTCCTTGAAAAACT  CCGCTCGAGGATAATTTTCAGCACTTGCT | 94 °C, 30 s; 58 °C, 30 s; 72 °C, 60 s; 35 cycles |
| STUB1-H260Q-F  STUB1-H260Q-R | TCGAGGAGCAGCTGCAGCGTG  CACGCTGCAGCTGCTCCTCGA | 94 °C, 30 s; 58 °C, 30 s; 72 °C, 90 s; 35 cycles |
| STUB1-Cherry-5’  STUB1-Cherry -3’ | CTAGCTAGCCACCATGAAGGGCAAGGAGG CCGCTCGAGTTAGTAGTCCTCCACCCAGCCA | 94 °C, 30 s; 58 °C, 30 s; 72 °C, 120 s; 35 cycles |
| **Primers for making shRNA** | | |
| sh-STUB1-1#-F sh-STUB1-1#-R | CCGGCCTTGTGCTACCTGAAGATTTCAAGAGAATCTTCAGGTAGCACAAGGCCTTTTTG  CTAGCAAAAAGGCCTTGTGCTACCTGAAGATTCTCTTGAAATCTTCAGGTAGCACAAGG | |
| sh-STUB1-2#-F  sh-STUB1-2#-R | CCGGAAGCGCTGGAACAGCATTGATTCAAGAGATCAATGCTGTTCCAGCGCTTCTTTTT  CTAGAAAAAGAAGCGCTGGAACAGCATTGATCTCTTGAATCAATGCTGTTCCAGCGCTT | |
| shYTHDF1-1#-F  shYTHDF1-1#-R | CCGGCACTGACTGGTGTCCTTTCTTTCAAGAGAAGAAAGGACACCAGTCAGTGCTTTTT  CTAGAAAAAGCACTGACTGGTGTCCTTTCTTCTCTTGAAAGAAAGGACACCAGTCAGTG | |
| shYTHDF1-2#-F shYTHDF1-2#-R | CCGGGTCTGGTCTCAGGACAAGTTTCAAGAGAACTTGTCCTGAGACCAGACCCTTTTT  CTAGAAAAAGGGTCTGGTCTCAGGACAAGTTCTCTTGAAACTTGTCCTGAGACCAGAC | |
| **Primers for real-time quantitative PCR analysis** | | |
| GAPDH-qF  GAPDH-qR | GTGGAGTCGGTCCTGTTCTATG  GTTACGCATGATCTGCACGAG | 95°C, 15s; 60°C, 15s; 72°C, 45s; 40 cycles |
| STUB1-qF  STUB1-qR | TCAAGGAGCAGGGCAATCGTCT  GCATCTTCAGGTAGCACAAGGC | 95°C, 15s; 60°C, 15s; 72°C, 45s; 40 cycles |
